# High-Throughput Mechanomic Screening Reveals Novel Regulators of Single-Cell Mechanics

**DOI:** 10.1101/2025.03.16.643502

**Authors:** Laura Strampe, Katarzyna Plak, Christine Schweitzer, Cornelia Liebers, Paul Müller, Marta Urbanska, Martin Kräter, Buzz Baum, Jona Kayser, Jochen Guck

## Abstract

The mechanical properties of cells are dynamic, allowing them to adjust to different needs in different biological contexts. In recent years, advanced biophysical techniques have enabled the rapid, high-throughput assessment of single-cell mechanics, providing new insights into the regulation of the mechanical cell phenotype. However, the molecular mechanisms by which cells maintain and regulate their mechanical properties remain poorly understood. Here, we present a genome-scale RNA interference (RNAi) screen investigating the roles of kinase and phosphatase genes in regulating single-cell mechanics using Real-Time Fluorescence and Deformability Cytometry (RT-FDC). Our screen identified 80 known and novel mechanical regulators across diverse cellular functions from 214 targeted genes, leveraging RT-FDC’s unique capabilities for comprehensive, high-throughput mechanical phenotyping with single-cell and cell cycle resolution. These findings refine our understanding of how signaling pathways coordinate structural determinants of cell mechanical phenotypes and provide a starting point for uncovering new molecular targets involved in biomechanical regulation across diverse biological systems.

**SIGNIFICANCE:** Cell mechanical properties are tightly regulated and play pivotal roles in processes ranging from tissue morphogenesis to disease progression. Despite their importance, the genetic regulation of single-cell mechanics remains largely unexplored. This study represents one of only a few large-scale mechanomic investigations conducted to date. It is the first study to leverage RT-FDC’s unique capability for high-throughput mechanical phenotyping with single-cell and cell cycle resolution to detect gene impacts that may be overlooked in lower-throughput or population-level studies. The mechanical genes identified here provide valuable data points for understanding how cells control their mechanical state and serve as a foundation for future studies exploring the molecular basis of biomechanical regulation.

## INTRODUCTION

Cell mechanics encompasses all physical interactions between a cell and its surroundings, including how cells generate, sense, and respond to mechanical forces. These mechanical properties are fundamental to numerous physiological processes, such as migration, proliferation, and differentiation (1–3), and are increasingly recognized as critical factors in pathological conditions, including cancer progression and immune responses. One key aspect of cell mechanics is cell stiffness, which describes how much a single cell deforms under external forces. Stiffer cells resist deformation, while softer cells deform more readily. Variations in cell stiffness have been implicated as biomarkers of diverse physiological and pathological states.

For instance, during chemically induced osteoblast differentiation of mesenchymal stem cells, cell softening outperformed traditional molecular biomarkers as a reliable indicator of differentiation (4). Red blood cells stiffen in pathological conditions such as sickle cell anemia, spherocytosis, malaria, and COVID-19 infection (5–8). In malaria, increased red blood cell stiffness elevates the risk of microcapillary obstruction and correlates with fatal patient outcomes (9). A more dynamic response is observed in neutrophils during immune response: they initially stiffen upon activation but soften significantly within 15 minutes, potentially aiding their migration towards the inflammation site (10). Similarly, single metastatic cancer cells are consistently softer than their healthy counterparts (11–13), in contrast to tumors, which tend to be stiffer than the surrounding healthy tissue as a whole (14). This softening in metastatic cancer cells is hypothesized to improve their ability to navigate dense tissue environments, enabling invasive behaviors such as tissue infiltration and intra- and extravasation (15). Another possibility is immune evasion, as softer cancer cells are not as readily killed by cytotoxic T-cells (16).

Despite extensive evidence correlating cell stiffness with disease states, the genetic and functional determinants of mechanical phenotypes remain poorly understood. Addressing this knowledge gap is the primary goal of mechanomics, a relatively new subfield of cell mechanics that attempts to connect the mechanical phenotype of cells to a comprehensive understanding of the underlying genotype. Current knowledge identifies the contractile actomyosin cortex as the primary regulator of cell stiffness. This dynamic network of actin filaments generates contractile forces through myosin motor proteins, with its activity tightly controlled by Rho signaling (17). Additional structural determinants of cell mechanics include other cytoskeletal proteins such as microtubules, intermediate filaments and septins, the lipid plasma membrane, the nucleus, and macromolecular packing in the cytoplasm (15). While cytoskeletal drugs can modulate cell stiffness by disrupting the different cytoskeletal filaments, they often induce cytotoxic effects (18, 19), limiting their utility in functional studies. Genetic approaches such as RNA interference (RNAi) could provide a more precise and less invasive means of modulating cell mechanics. RNAi achieves gene knockdown by introducing short RNA strands that bind to corresponding mRNA transcripts, preventing their translation into proteins (20). Large-scale RNAi screens systematically evaluate the effects of individual gene knockdowns, enabling the precise identification of key genetic regulators of cell stiffness. Previous mechanomic RNAi screens using atomic force microscopy (AFM) identified both known and novel regulators of actin organization and cortical stiffness (21, 22). However, the low throughput of AFM limited the number of cells analyzed per gene, raising concerns about statistical power and reproducibility. Alternatively, transcriptomics-based mechanomics has been used to correlate gene expression with mechanical phenotypes, offering insights into the collective effects of many genes, though it lacks the precision of targeted knockdown approaches (23, 24). The common fruit fly *Drosophila melanogaster* serves as an ideal model organism for RNAi studies due to its relatively simple genome, conservation of many human homologs, and the efficiency of RNA strand uptake (25, 26).

Real-Time Fluorescence and Deformability Cytometry (RT-FDC) enables high-throughput measurement of cell mechanics at the single-cell level (27). Cells are suspended in a viscous measurement buffer and subjected to hydrodynamic shear forces as they flow through a narrow microfluidic channel. Through high-speed image analysis, RT-FDC measures how much each cell deforms, from which its apparent Young’s modulus can be estimated (28). This method can analyze up to 100 cells per second, enabling the rapid mechanical profiling of entire cell populations while retaining single-cell resolution (29). In addition, RT-FDC supports subcellular localization of fluorescent proteins, enabling the differentiation of interphase and mitotic cells by detecting nuclear envelope breakdown based on nuclear and cytoplasmic fluorescent peak widths (30). This allows for the separate mechanical phenotyping of both cell cycle phases in phase-mixed cell populations.

The feasibility of RT-FDC for mechanomic screening was first demonstrated by Rosendahl et al. (30) through a targeted RNAi screen of 42 genes associated with actin cortex regulation. This study confirmed the critical role of Rho signaling in controlling cell stiffness but remained limited to a small gene set previously known to influence the actin cytoskeleton. RT-FDC was subsequently applied in a clinical context for biophysical phenotyping of blood cells in Chorea-Acanthocytosis patients (31) and circulating immune cells in systemic sclerosis patients (32). In this work, we significantly expand the scope of RT-FDC-based mechanomic screening by targeting 214 kinase and phosphatase genes in *Drosophila melanogaster*. Kinases and phosphatases modulate protein activity through reversible phosphorylation, inducing conformational changes that switch proteins between active and inactive states and thus regulating numerous downstream signaling pathways (33). Among both enzyme types, phosphatases are generally less well studied, making them ideal candidates for identifying novel genetic regulators of cell mechanics (34). Our study not only identifies potential new regulators of cell stiffness but also emphasizes the distinct advantages of incorporating cell cycle resolution into mechanomic screening. We demonstrate that resolving mechanical phenotypes at the single-cell and cell cycle level uncovers phase-specific gene functions, prevents the identification of apparent stiffening effects caused by shifts in cell cycle distribution, and improves the accuracy and reliability of mechanical hit detection.

## MATERIALS AND METHODS

### Cell culture

We used the *KC167* cell line (*Drosophila melanogaster*) for its high compatibility with RNAi and its uniform cell size and spherical morphology (35), making it particularly suitable for RT-FDC measurements, which assume cells to be spherical prior to deformation inside the channel (36). Additionally, the hemocyte-like *KC167* cells naturally grow in suspension (37), meaning they are measured in their native state by RT-FDC. The semi-adherent *KC167* cell line, stably expressing nuclear NLS-GFP and cytoplasmic tdTomato (30), was cultured in Shield and Sang M3 Insect Medium (Sigma Aldrich, Darmstadt, Germany, S3652) supplemented with 10 % Fetal Bovine Serum (FBS, Sigma Aldrich, Darmstadt, Germany, F7524) and 1 % Penicillin-Streptomycin (Pen-Strep, Thermo Fisher, Darmstadt, Germany, 15070063). The FBS was heat-inactivated for 30 minutes at 56 °C. This complete growth medium was stored at room temperature for up to two weeks. The cells were cultured in T25 cell culture flasks (Greiner Bio-One, Frickenhausen, Germany, 690175) at 24 °C without CO_2_. Growth medium was exchanged every other day. Cells were passaged once a week. For passaging, the old culture medium was removed and the cells were resuspended in 2 ml of fresh complete growth medium by repetitive pipetting. 300 µl of this cell suspension was then added to a new T25 flask containing 4 ml of complete growth medium.

### Gene knockdown via RNA interference

RNA interference (RNAi) was induced in the *KC167* cells via transfection with double-stranded RNA (dsRNA). We used a subset of the Heidelberg 2 dsRNA library (HD2 or BKN library) (38), targeting 143 phosphatase genes and 71 kinase genes (see Table S1). This gene set covers the majority of the *Drosophila* phosphatome, along with several well-characterized kinases facilitating comparison with previous mechanical studies. The library contains two different dsRNA strands for each target gene. Sequence information, knockdown efficiency and siRNA specificity for all strands can be accessed via the RNAi identifier under http://www.genomernai.org/.

Cells were harvested for dsRNA transfection at 60% to 80% confluency. They were removed from the culture flask and resuspended in fresh medium (see passaging protocol described above) and then centrifuged at 100 ×g and room temperature for 5 minutes to form a cell pellet. The cells were then washed with 3 ml of Dulbecco’s phosphate-buffered saline (DPBS, Thermo Fisher, Darmstadt, Germany, 14190250), which had been adjusted to a pH of 6.8 and an osmolality of 320 mOsm/kg and is hereafter referred to as Drosophila-PBS. pH was adjusted through addition of NaOH or HCl, osmolality through addition of NaCl. After washing, the cells were centrifuged again as described above and resuspended in 500 µl serum-free medium (Shield and Sang M3 Insect Medium supplemented with 1 % Pen-Strep). The cells were then counted using the Countess II Automated Cell Counter (Thermo Fisher, Darmstadt, Germany, AMQAX1000) by diluting 10 µl of the cell suspension in 40 µl of Drosophila-PBS and staining the diluted cells 1:1 with Trypan Blue (Thermo Fisher, Darmstadt, Germany, T10282). Based on this cell count, the cell suspension was further diluted with serum-free medium to reach a final concentration of 5×10^6^ cells/ml.

For each knockdown sample, 2.7 µl of dsRNA from the BKN library (1 µg/µl stock solution) was mixed with 45 µl serum-free cell suspension. For the control samples, 2.7 µl of Tris-EDTA buffer (Sigma Aldrich, Darmstadt, Germany, 93283) was used instead. The cell-dsRNA mix was transferred to a 24-well plate (Greiner Bio-One, Frickenhausen, Germany, 662160), where it was placed as a single droplet in the center of one well and left to settle for 1 hour. This well was then filled with 400 µl of complete growth medium and incubated for 4 days at 24 °C without CO_2_. After 2 days of incubation, 300 µl of additional complete growth medium was added to the well and the cells were resuspended by repetitive pipetting. This *Drosophila* RNAi protocol has been reliably applied in a range of previous studies (30, 39–41).

### Enrichment of the mitotic fraction

Given the typically small number of mitotic cells in asynchronous cell populations, we enriched the mitotic fraction of each cell sample through incubation with either 5 µM or 50 µM Colchicine (Thermo Fisher, Darmstadt, Germany, J61072.06) prior to the mechanical phenotyping measurement, depending on mitotic yield. For this, the Colchicine stock solution (100 mM in ethanol) was diluted at a 1:20 000 or 1:2 000 ratio in complete growth medium, respectively. After removal of the dsRNA-medium from the knockdown sample well, 300 µl of diluted Colchicine was added to the well and incubated in the dark for 5 hours. This treatment has been shown to increase the fraction of mitotic cells in *KC167* cultures from approximately 5 % to up to 17 % (30).

### Mechanical phenotyping using Real-Time Fluorescence and Deformability Cytometry (RT-FDC)

Mechanical measurements took place 4 days after RNAi knockdown treatment. The RT-FDC setup, measurement procedures and real-time image analysis protocols have been described in detail in (30). For this study, we used microfluidic chips with a microfluidic channel width of 20 µm and a total flow rate of 0.04 µl/s. Fluorescence traces were collected in the green 525/50 nm channel for nuclear NLS-GFP, and in the red 593/46 nm channel for cytoplasmic tdTomato. The viscous measurement buffer used for this study was prepared by supplementing Drosophila-PBS (described above) with 0.5% (w/v) methylcellulose (Fisher Scientific, Reinach, Switzerland, 11374247), hereafter referred to as MC. The MC was warmed to room temperature and sterile filtered through a 0.22 µm PVDF filter (Merck, Darmstadt, Germany, SLGV033RB) before measurements. Cells were collected and centrifuged into a cell pellet as described above, and then resuspened in MC and loaded onto the microfluidic chip for mechanical phenotyping as described by Rosendahl et al. (30). In total, at least 2000 single cells were measured for each cell sample.

All measured RT-FDC datasets are available on the Deformability Cytometry Open Repository (DCOR) under https://dcor.mpl.mpg.de/organization/mechano-genomic-screening. DCOR is an effort towards making Deformability Cytometry (DC) data easily accessible to anyone. DCOR serves DC datasets (normally around 25 GB in size) for download or for online data analysis using the graphical user interface Shape-Out (https://shapeout.readthedocs.io). In addition, DCOR facilitates sharing and organizing DC data online. The public DCOR instance at https://dcor.mpl.mpg.de is free to use for the general public. For private projects, it is also possible to self-host DCOR. More information can be found at https://dc.readthedocs.io.

Post-experimental data analysis was performed using the python package dclab version 0.62.6 (available at https://github.com/DC-analysis/dclab). Measured cell events were filtered based on features extracted from the bright-field images and fluorescence traces, explained in detail in Herbig et al. (42), retaining only events within the following parameter ranges: cross-sectional cell area between 20 and 200 µm^2^, porosity (also called area ratio) between 1 and 1.05 and maximum fluorescence intensities in both channels between 400 and 40000 intensity units. In addition, manual gating in a plot of cell position within the bright-field image against peak position of the fluorescence trace was used to filter out events were bright-field image and fluorescence trace did not originate from the same cell. Manual gating was also used to separate the interphase and mitotic subpopulations within each sample based on peak widths in the two fluorescence channels (compare Fig. 1A). Young’s modulus values were calculated based on a cell’s area and deformation using the lookup table LE-2D-FEM-19 implemented in dclab.

**Figure 1:**
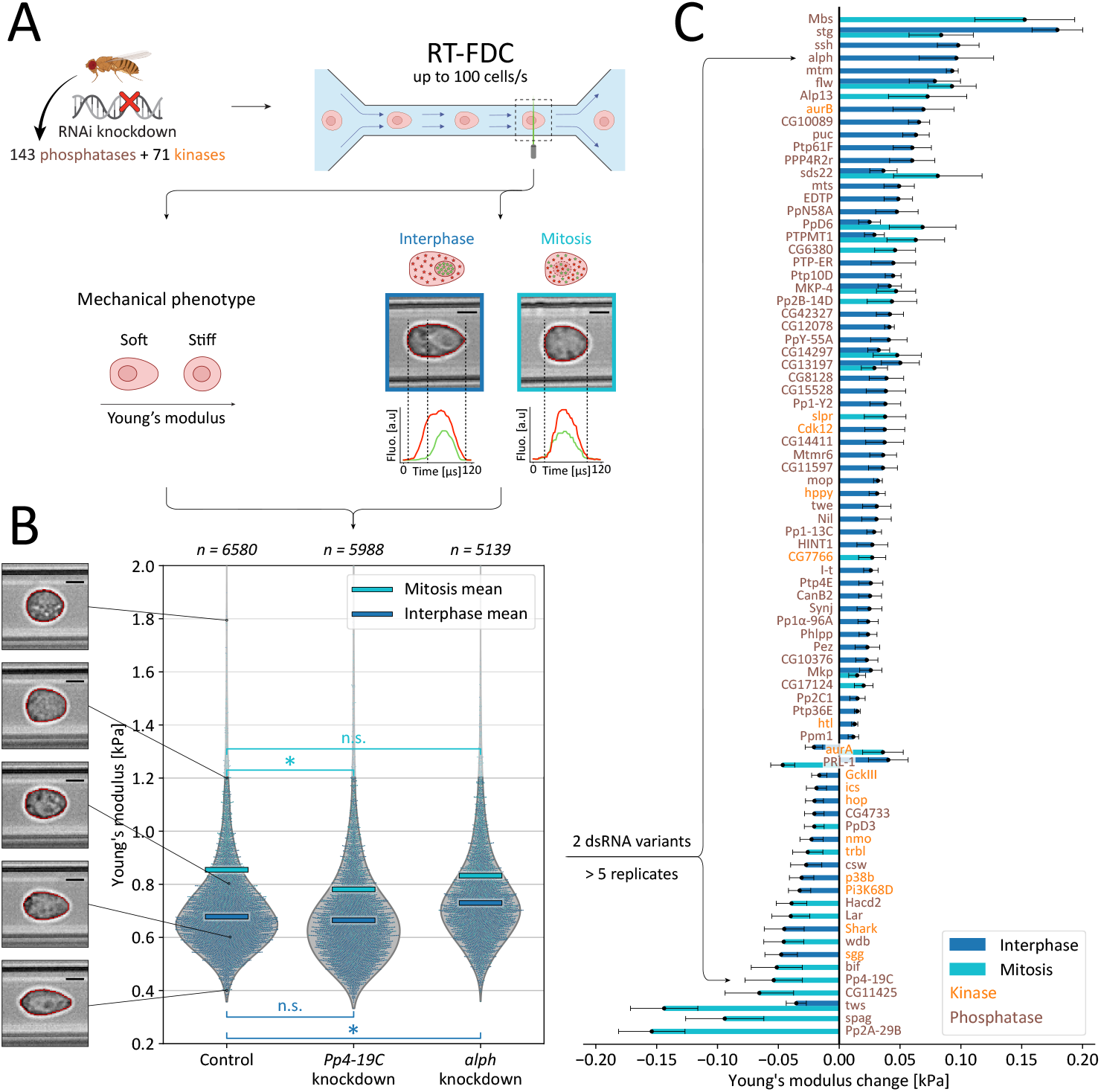
Mechanical knockdown screen of 214 kinase and phosphatase genes. **A** Overview of the mechanomic screening workflow. For each target gene, RNAi knockdown was performed in the *Drosophila* cell line *KC167*. Real-Time Fluorescence and Deformability Cytometry (RT-FDC) was used to determine the stiffness and cell cycle phase of at least 2000 single cells for each knockdown sample. Cell stiffness was quantified as Young’s modulus based on the observed hydrodynamic deformation, while cell cycle phase was inferred from fluorescent labeling of nuclear envelope breakdown. **B** Typical Young’s modulus distributions for cell cycle specific knockdown effects. Each dot represent a single cell measurement, blue for interphase and cyan for mitotic cells. Horizontal lines indicate the mean Young’s modulus for each cell cycle phase. Gray density plots represent the full population distrubtion without cell cycle-resolution. *Pp4-19C* knockdown causes softening of mitotic cells, *alph* knockdown causes stiffening of interphase cells. Five representative bright-field microscopy images of single cell measurements at different stiffness values are shown. Scale bars are 5 µm. For the control, *Pp4-19C* knockdown and *alph* knockdown samples, 4, 17 and 14 measurements are above the plot window going up to 3.6 kPa, 8.3 kPa and 3.0 kPa, respectively. **C** Knockdown effects of all mechanical hits (80/214). For each target gene, at least five biological replicates were measured using two distinct dsRNA strands. Genes causing a significant (*p < 0*.*05*) change in mean Young’s modulus are regarded as mechanical hits. Positive Young’s modulus changes indicate cell stiffening, negative changes indicate cell softening upon gene knockdown. Phosphatase genes are labeled in brown, kinase genes in orange. Mean difference calculation, error margin estimation and significance testing were performed through linear mixed model analysis. Residuals were not normally distributed (Shapiro–Wilk, mean of all measurements, *W = 0*.*76, p < 0*.*001*).

### Screening procedures and statistical analysis

For each target gene, at least 5 biological replicates using two different dsRNA strands were measured. Genes were divided into three groups (*kin* for all kinases and *phos1*/*phos2* for phosphatases) which were measured consecutively. For experimental blinding, all dsRNA strands, including those targeting the same gene, were assigned unique sample IDs within their group (see Table S1). Replicates of the same sample ID were measured on different days, with several weeks between consecutive measurements. On each measurement day, around 10 different gene knockdowns and at least 2 independent control samples were measured, with the first measurement of the day always being a control. The same microfluidic chip was used for all measurements on a given day, except for when clogging in the microfluidic channel necessitated its replacement. In that case, an additional control sample was measured directly after chip exchange. This rigorous scheme of temporally distributed replicates and multiple daily controls was implemented to take unavoidable day-to-day variation, such as changes in ambient temperature, MC buffer viscosity or Colchicine concentration, into account. Between measurements, 50 µl of MC was loaded onto the chip to separate samples as described in Rosendahl et al. (30). During analysis, each knockdown measurement was matched to the temporally closest control measurement which had used the same chip.

Statistical analysis was performed using a linear mixed model (LMM) to estimate mean differences in Young’s modulus distributions between knockdown samples and matched daily controls. This model accounts for baseline variability in Young’s modulus values, caused by changes in environmental conditions or knockdown efficiency across experimental days. Significance was determined using a likelihood ratio test, which is robust against the large sample sizes typical of RT-FDC datasets. It should be noted that LMM analysis assumes normally distributed data, a condition that is typically not met in RT-FDC datasets. Nonetheless, LMM analysis has proven to be a robust and widely adopted approach for analyzing deformability cytometry data (23, 30, 31, 43, 44). A detailed discussion of the use of linear mixed models for deformability cytometry data is provided by Herbig et al. (36). A gene was identified as a mechanical hit, if mean Young’s modulus differed between knockdown samples and controls with *P* < 0.05. For the mitotic downsampling reanalysis, mean differences and p-values were averaged over 10 downsampling repetitions to account for stochastic variability. Graphs were generated using the python packages matplotlib and seaborn. Schematics were created in BioRender and Adobe Illustrator. BioRender schematics can be accessed via https://BioRender.com/f63v496.

## RESULTS

### Population-wide single-cell mechanics screening using RT-FDC

We systematically investigated how genetic perturbations influence single-cell mechanics by performing a genome-scale RNA interference (RNAi) knockdown screen (Fig. 1A) targeting the entire *Drosophila* phosphatome (143 phosphatase genes) and 71 additional kinase genes with mostly known gene function using the Heidelberg 2 (BKN) RNAi library (38).To quantify changes in cell mechanics upon gene knockdown, we used Real-Time Fluorescence and Deformability Cytometry (RT-FDC), a high-throughput microfluidic mechanical phenotyping technique. In short, RT-FDC quantifies the apparent Young’s modulus of single cells in suspension based on their hydrodynamic deformation in a narrow flow channel at up to 100 cells/s. In addition, it distinguishes interphase and mitotic cells based on fluorescent labeling (Fig. 1A, see Introduction and Methods for details). This enables us to account for the well-established mechanical changes during cell cycle progression, specifically the increased stiffness of mitotic cells (30, 45, 46), by separating the interphase and mitotic subpopulations during statistical analysis (Fig. 1B).

Among the 214 targeted genes, we identified 80 mechanical hits that significantly shifted the mean Young’s modulus of the knockdown cell samples compared to matched daily controls in at least one cell cycle phase (Fig. 1C). The measured knockdown effects had a median magnitude of 0.04 kPa, compared to a typical standard deviation of 0.24 kPa within one sample. The majority of these mechanical hits (57 genes) caused cell stiffening upon knockdown, 21 caused softening and two gene knockdowns (*aurA* and *PRL-1*) had opposing effects on interphase and mitotic cells. The strongest stiffening effects were observed in interphase *stg* knockdown cells and mitotic *Mbs* knockdown cells, which showed a 0.18 kPa and 0.15 kPa increase in stiffness, respectively. Knockdown of *stg*, a known regulator of mitotic entry in early *Drosophila* embryos (47), also stiffened mitotic cells to a smaller extend (0.08 kPa). *Mbs* (*Myosin binding subunit*) is a subunit of the dimeric *Drosophila* myosin light chain phosphatase, which dephosphorylates the regulatory light chain of non-muscle myosin II to promote post-contractile relaxation (48, 49). Knockdown of *flw*, another subunit of the complex, also stiffens both interphase and mitotic cells, while direct knockdown of the light chain encoding gene *sqh* has previously been shown to soften mitotic cells (30), suggesting that loss of the light chain impairs contraction, whereas loss of the phosphatase locks cells in a contracted, stiffened state. The strongest softening effects were caused by *Pp2A-29B, spag* and *tws* knockdown in mitotic cells (0.15 kPa, 0.09 kPa and 0.14 kPa decrease in Young’s modulus), with *tws* knockdown also weakly softening interphase cells by 0.035 kPa. *Spag*, a homolog of human *RPAP3*, is involved in diverse cellular processes, including the assembly of cellular machineries (50). *Pp2A-29B* and *tws* encode subunits of the *Drosophila* Protein Phosphatase 2A (PP2A) complex, a multifunctional regulator of cell cycle progression, morphology, and development (51). Additional PP2A family subunits identified as hits in our screen include *mts* and *wdb*.

### Comparison to full population analysis

Among the identified genes, *PRL-1* and *aurA* stand out by exhibiting opposing mechanical effects between the cell cycle phases. Knockdown of the phosphatase *PRL-1*, a known suppressor of the *SRC* oncogene and marker of cancer metastasis (52), stiffened interphase cells while softening mitotic cells, whereas knockdown of *aurA*, a mitotic kinase known for its roles in regulating mitotic entry, centrosome separation, and spindle formation (53), softened interphase cells and stiffened mitotic cells. These bidirectional effects raise the question of how the knockdown of these genes would have been recorded in a standard screening protocol which does not account for cell cycle progression. To investigate this, we conduct a full population analysis, where interphase and mitotic cells are pooled into a single sample for each gene knockdown instead of being separated (gray density plots in Fig. 1B). We then compare the mechanical hits identified in our cell cycle-resolved analysis to the effects observed in this full population analysis (Fig. 2).

**Figure 2:**
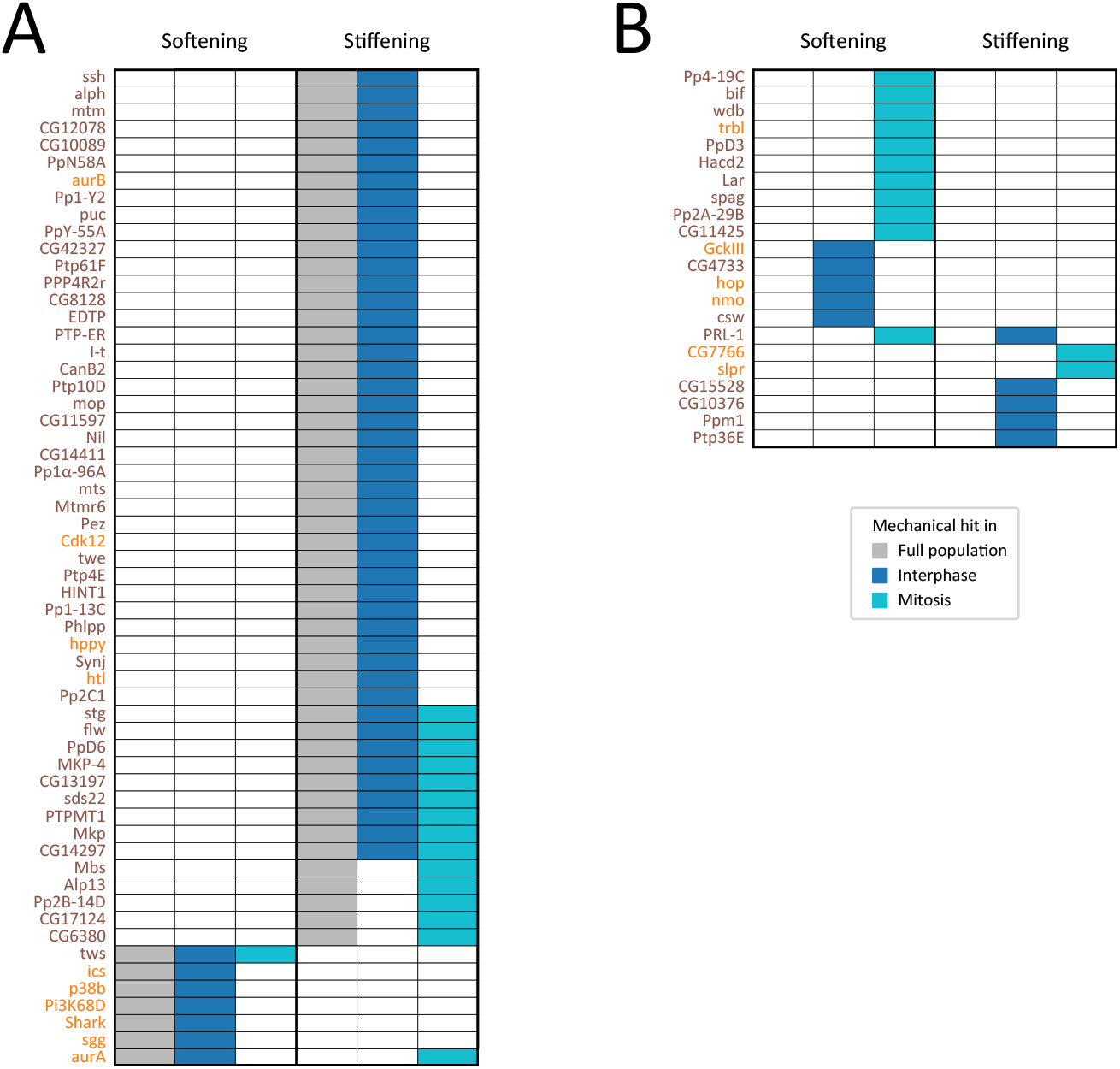
Comparison of mechanical hits identified from the cell cycle-resolved analysis, separately comparing mean Young’s modulus values in the interphase (blue) and mitotic (cyan) subpopulations (see Fig. 1), and from the full population analysis (gray). Genes in **A** are detected as hits by both analyses, while genes in **B** appear exclusively in the cell cycle-resolved analysis. Hits are separated into softening (Young’s modulus change < 0) and stiffening (Young’s modulus change > 0) knockdown phenotypes. Phosphatase genes are labeled in brown, kinase genes in orange. Significance testing was performed through linear mixed model analysis.

Of the 80 genes originally identified as mechanical hits in the cell cycle-resolved analysis, 58 can also be detected in the full population analysis (Fig. 2A). Among these, 11 genes influenced both interphase and mitotic cells, 42 genes affected only interphase cells and just five genes exclusively changed the mechanical phenotype of mitotic cells. Most full population hits cause cell stiffening upon gene knockdown. This trend is particularly pronounced in the 16 full population hits that also influence mitotic mechanics, with only one gene knockdown causing cell softening (*tws*), while all other caused cell stiffening.

22 genes identified as mechanical hits in the cell cycle-resolved analysis did not show any significant effect in the full population analysis (Fig. 2B). The majority of these hidden mechanical hits induced cell softening upon knockdown, contrasting the predominance of stiffening phenotypes in the full population analysis. 10 gene knockdowns exclusively softened mitotic cells, a phenomenon entirely absent from the full population results (Fig. 2A). There, the only mitotic softening knockdown was *tws*, which also softened interphase cells. In total, the cell cycle-resolved analysis more than doubled the number of discovered cell softening phenotypes compared to a full population analysis.

Notably, the full population analysis returned 12 additional stiffening phenotypes that were not detected in either the interphase or mitotic subpopulations. These apparent knockdown effects can be attributed to two factors: drug-induced enrichment of the inherently stiffer mitotic fraction, and the increased statistical power of the larger full population. A detailed discussion of these apparent mechanical phenotypes is provided in the Supplemental Material.

### Known and novel regulators in diverse cellular functions

To evaluate the impact of our screen, we classified the 80 mechanical hits identified among the 214 screened kinase and phosphatase genes into three categories based on literature: known mechanical regulators, mitotic regulators, and novel candidate genes without prior links to cell mechanics or mitosis (Fig. 3).

**Figure 3:**
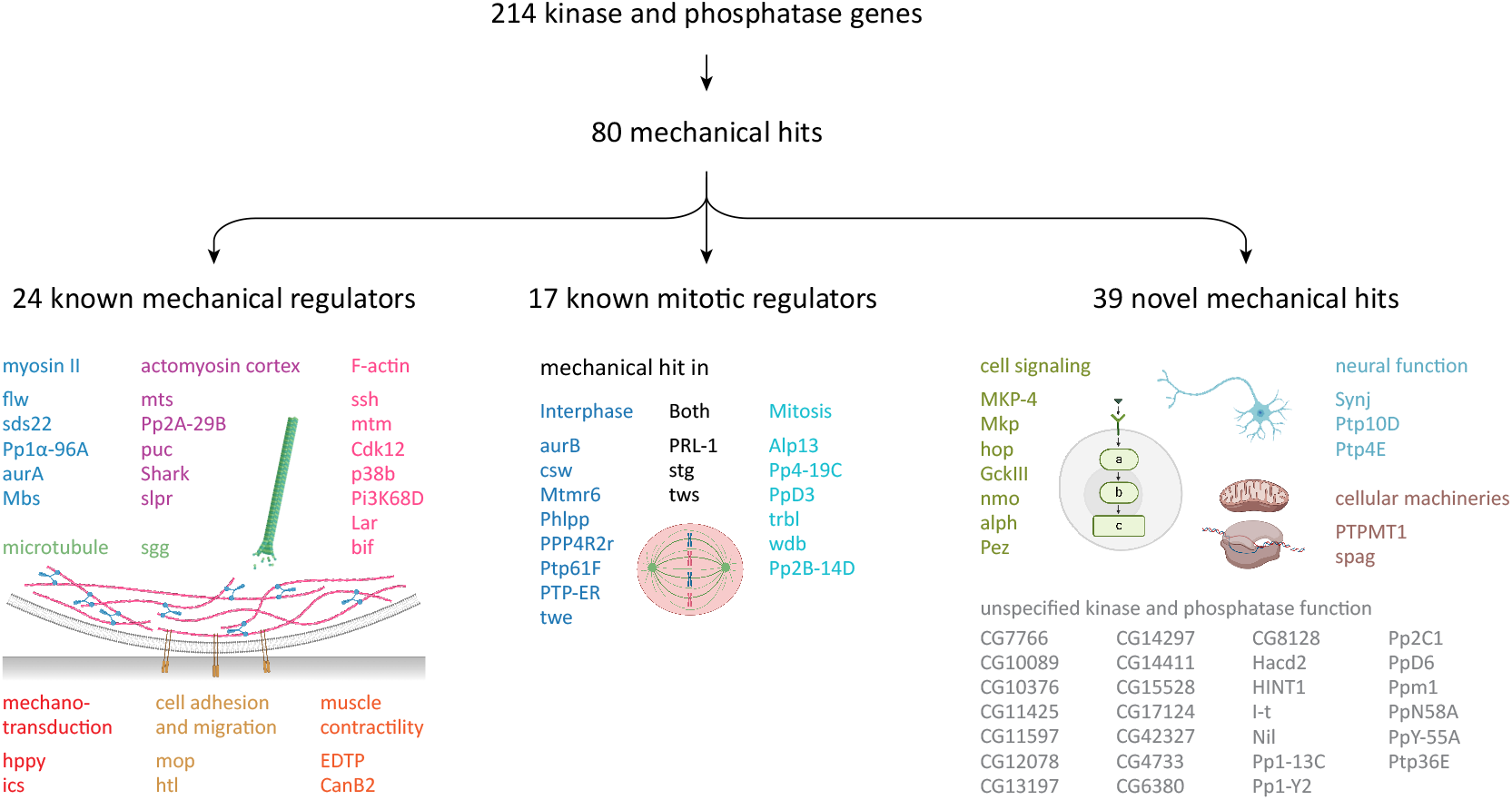
Classification by gene function of the 80 mechanical hits identified in our mechanomic screen according to literature. Mechanical regulators only implicated in regulation of mitotic mechanics are classified together with regulators of cell cycle progression through mitosis as mitotic regulators. Genes neither known for mechanical nor mitotic regulation are classified as novel mechanical hits. For known mechanical regulators and novel genes, colors indicate subclasses of cell function. For all subclasses except actomyosin cortex, mechanotransduction, muscle contraction and unspecified kinase and phosphatase function, text colors correspond to component colors used in the schematic drawings. For known mitotic hits, colors indicate the cell cycle in which the mechanical effects identified in this screen were identified. Literature sources in Tables 1,2 and 3.

24 genes were classified as known mechanical regulators, validating RT-FDC as a robust screening platform. Most of these genes are involved in actomyosin regulation, either through regulation of cortical tension via JNK signaling (*puc, Shark, slpr*) or as components of the PP2A complex (*Pp2A-29B, mts*). Others contribute to F-actin filamentation and organization (*ssh, mtm, Cdk12, p38b, Pi3K68D, Lar, bif*) or the phosphorylation of the non-muscle myosin II regulatory light chain (*Mbs, flw, PP1β9c, sds22, AurA*). Beyond actomyosin regulation, the screen identified genes linked to diverse mechanical cell functions. Some play roles in F-actin organization during adhesion and migration (*mop, htl*), while others contribute to mechanotransduction (*hppy, ics*) or indirect flight muscle (IFM) contractility in *Drosophila* (*EDTP, CanB2*). Regulation of the microtubule cytoskeleton was also represented (*sgg*). The identification of genes spanning multiple known mechanics-related pathways highlights the versatility of RT-FDC-based mechanical phenotyping, demonstrating its potential for systematically mapping genetic regulators of cell mechanics.

**Table 1:**
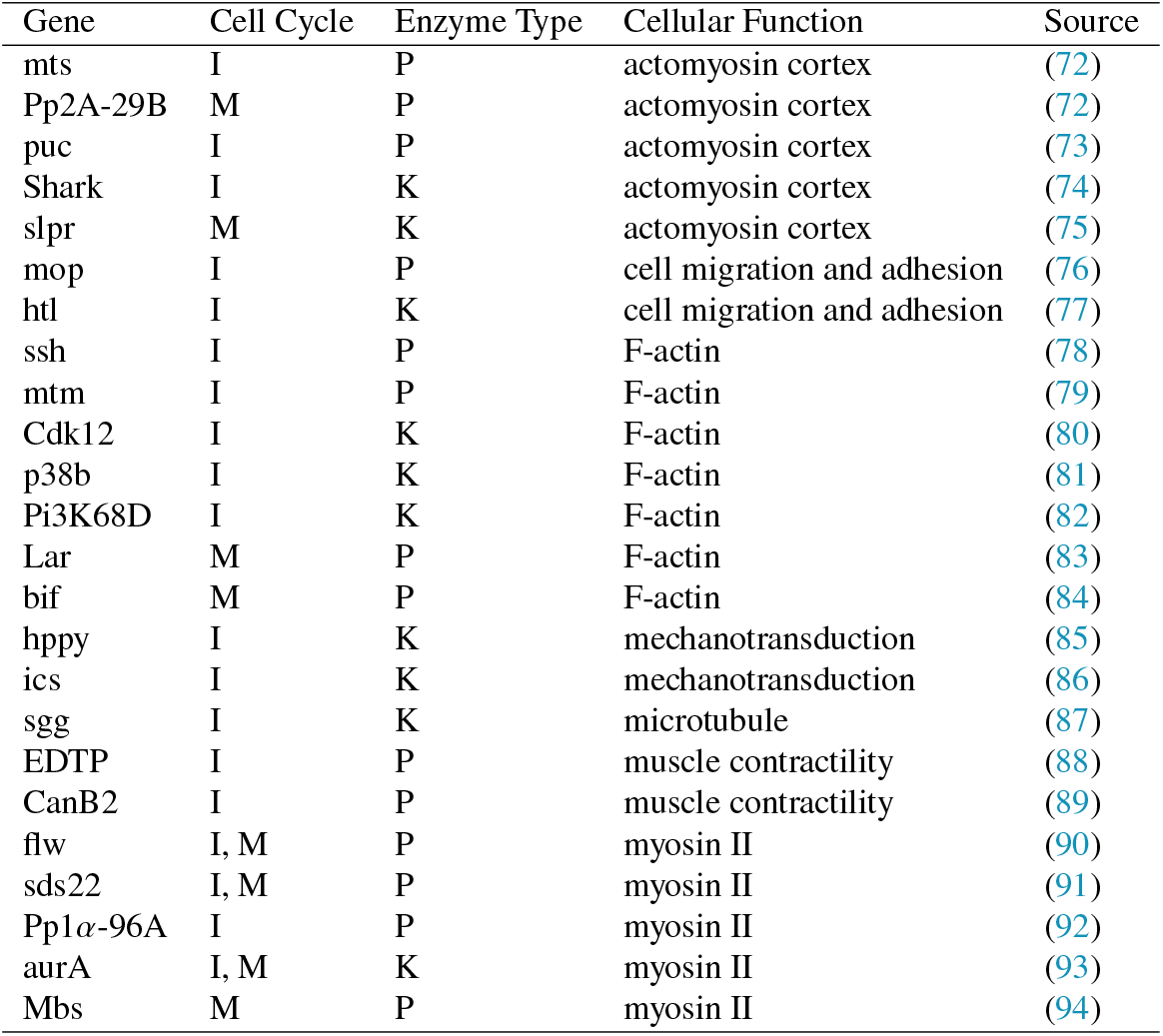
Classification of gene function for known regulators of cell mechanics identified by the mechanomic screen according to literature. Genes only implicated in mitotic mechanics are instead classified as mitotic regulators. I/M for interphase- or mitosis-specific gene knockdown effects. P/K for phosphatase or kinase gene.

**Table 2:**
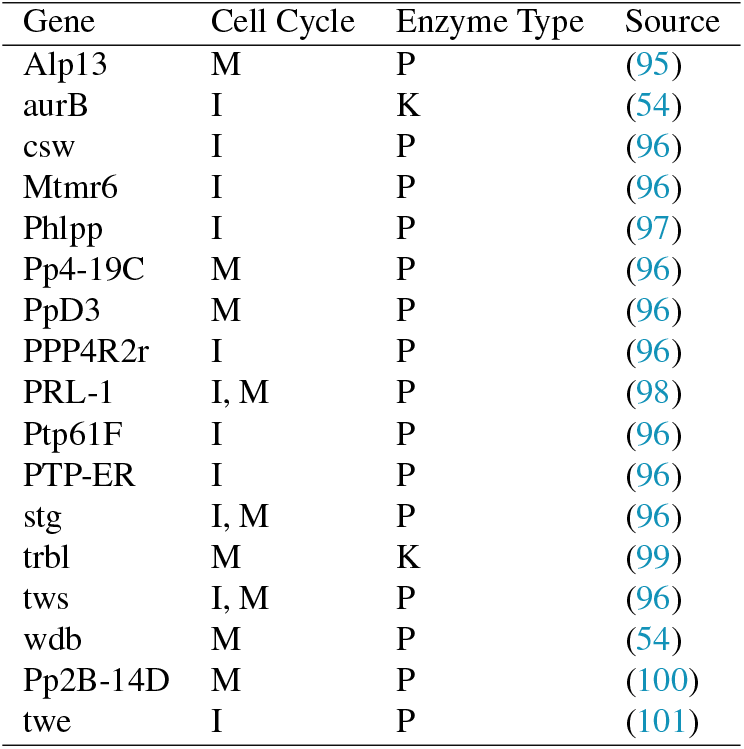
Known regulators of mitosis or meiosis identified by the mechanomic screen according to literature. I/M for interphase- or mitosis-specific gene knockdown effects. P/K for phosphatase or kinase gene.

**Table 3:** Classification of gene function for genes neither implicated in mechanical nor mitotic regulation identified by the cell cycle-resolved mechanomic screen according to literature. I/M for interphase- or mitosis-specific gene knockdown effects. P/K for phosphatase or kinase gene. Genes whose function is not well established beyond their general kinase or phosphatase function are not listed.

| Gene | Cell Cycle | Enzyme Type | Cellular Function | Source |
| --- | --- | --- | --- | --- |
| Synj | I | P | neural function | (102) |
| Ptp10D | I | P | neural function | (103) |
| Ptp4E | I | P | neural function | (103) |
| MKP-4 | I,M | P | signal transduction | (104) |
| Mkp | I,M | P | signal transduction | (105) |
| hop | I | K | signal transduction | (106) |
| GckIII | I | K | signal transduction | (107) |
| nmo | I | K | signal transduction | (108) |
| alph | I | P | signal transduction | (109) |
| Pez | I | P | signal transduction | (110) |
| spag | M | P | cellular machineries | (50) |
| PTPMT1 | I,M | P | cellular machineries | (111) |

Several mechanical hits were previously associated with microtubule dynamics during mitotic spindle assembly, including *wrd, aurB*, and *Pp4-19C* (54, 55). These genes were not included in the known mechanical regulators category but were instead grouped as mitotic regulators. Since mitosis inherently involves mechanical processes such as cytoskeletal reorganization and spindle assembly, most mitotic regulators were expected to affect cell stiffness, either directly or indirectly, particularly in mitotic cells. In total, we identified 17 known mitotic regulators whose knockdown significantly altered cell mechanics. Of these, nine genes changed mechanics of mitotic cells (*stg, Alp13, Pp2B-14D PpD3, trbl, PRL-1, Pp4-19C, tws, wdb*). Notably, three of theses mechanical hits in mitosis (*stg, PRL-1, tws*) and eight additional genes (*aurB, Ptp61F, PPP4R2r, PTP-ER, Mtmr6, twe, Phlpp, csw*) also cause a significant mechanical shift in the interphase cell population. This indicates that these genes have regulatory roles beyond mitosis, suggesting previously unrecognized gene functions in interphase cell mechanics. Finally, our screen identified 39 genes with no prior connection to either cell mechanics or mitosis. This set includes well-characterized genes involved in various signaling pathways (*MKP-4, Mkp, hop, GckIII, nmo, alph, Pez*), as well as genes linked to neural function (*Synj, Ptp10D, Ptp4E*) and genes involved in assembly and activity regulation of cellular machineries (*spag, PTPMT1*). All other mechanical hits in this category have not been characterized beyond their general kinase and phosphatase function. The identification of such diverse novel mechanical regulators underscores the power of RT-FDC for genome-scale mechanical screening in uncovering novel regulatory pathways and provides a foundation for future studies exploring the genetic basis of single-cell mechanics.

## DISCUSSION

Cells actively regulate their mechanical properties to adapt to changing biological contexts, yet the mechanistic links between genotype and single-cell mechanical phenotype remains poorly understood. In this study, we presented a genome-scale RNA interference (RNAi) knockdown screen of kinase and phosphatase impact on single-cell mechanics using Real-Time Fluorescence and Deformability Cytometry (RT-FDC).

### Cell cycle resolution improves mechanical screening

Our screen utilized the cell cycle resolution capabilities of RT-FDC to separately analyze the mechanical effect of gene knockdown on interphase and mitotic cells. We demonstrated that compared to the full population analysis, the cell cycle-resolved analysis identified additional mechanical hits, decreased bias by uncovering softening and mitosis-specific effects overlooked by full population analyses, and effectively eliminated apparent stiffening effects that were caused by knockdown-induced shifts in cell cycle distribution rather than genuine changes in single-cell mechanics.

One mechanical hits uniquely identified by the cell cycle-resolved analysis was *PRL-1*, which directly counteracted the mechanical oscillations associated with cell cycle progression by stiffening interphase cells and softening mitotic cells. This bidirectional mechanical shift upon *PRL-1* knockdown suggests that overexpression of *PRL-1* may amplify these mechanical oscillations, broadening the range of mechanical phenotypes a single cell can adopt. Given *PRL-1*’s established role as a cancer metastasis marker (52), such mechanical adaptability could facilitate tumor progression by enhancing a cell’s ability to overcome physical constraints, including detachment from the primary tumor, migration through dense tissues, and invasion into and out of the vasculature. This hypothesis aligns with findings from epithelial–mesenchymal transition (EMT), a key process in cancer metastasis, where breast epithelial cells were shown to stiffen in mitosis and soften in interphase, increasing single-cell mechanical variability (56). The discovery of *PRL-1* therefore highlights the strength of cell cycle resolution in uncovering biologically relevant gene functions that remain obscured in population-wide analyses. It should be noted that Colchicine, used to enrich the mitotic fraction in this study, disrupts microtubule dynamics. This effect, included in our controls, has been shown to influence cell mechanics (57–59).

### Comparison with previous mechanomic screens

The genome-scale mechanical screen presented in this work significantly expands the scope of deformability cytometry-based screens, surpassing prior studies which were limited to smaller target groups, such as 42 cytoskeletal regulator genes (30) or a limited number of chemical compounds (60). In contrast, our study evaluated a comprehensive set of 143 phosphatase genes and an additional 71 kinase genes in *Drosophila*, totaling 214 target genes. By targeting the entire *Drosophila* phosphatome, it represents an important step toward establishing RT-FDC as a viable platform for genome-scale mechanomic screening. However, a key limitation of RT-FDC lies in its inherently sequential measurement process: unlike typical high-throughput assays that can process many samples in parallel, RT-FDC analyzes one sample at a time. As a result, large-scale screens require the sequential measurement of hundreds of individual samples, limiting overall throughput and creating a practical bottleneck for comprehensive, full-genome screens involving thousands of conditions. Overcoming this limitation will require future advances in parallelization, enabling simultaneous measurement of multiple samples, or process automation, allowing continuous, unattended operation.

Larger mechanomic screens have previously been performed using different mechanical phenotyping methods. Toyoda et al. (22) conducted a RNAi-based screen of over 1,000 human genes using atomic force microscopy (AFM), including targets involved in the actomyosin cortex, membrane-associated transporters, and kinases, specifically analyzing the mechanical properties of mitotic cells. However, the low throughput of AFM limited their measurements to only a few cells per knockdown condition, compared RT-FDC’s high-throughput capacity of thousands of cells per condition, increasing their screens susceptibility to sampling bias. Our results demonstrate that capturing the full distribution of mechanical phenotypes is critical, as the knockdown-induced shifts in cell stiffness were typically smaller than the inherent biological variability observed within the cell populations. Han et al. (61) employed a pooled CRISPR-Cas9-mediated loss-of-function screen targeting 507 human kinase genes using microfluidic sorting to identify gene knockouts overrepresented among softer mechanical phenotypes. While this approach significantly increased throughput compared to AFM, it lacked single-cell resolution, providing no detailed stiffness distributions for individual gene knockouts. Thus, RT-FDC uniquely combines high throughput with single-cell resolution, addressing the limitations of both methods.

By comparing our results to these previous studies via cross-species homology, we observed partial overlap in identified hits. Of the 71 kinases targeted in our study, 66 were also tested by Toyoda et al. (22), identifying 11 as mechanical hits; two of these hits (*hppy, hop*) were replicated in our study. Similarly, Han et al. (61) tested 54 kinases targeted in our screen, identifying seven hits, five of which overlapped with ours (*p38b, hop, sgg, aurA, slpr*). This cross-species validation underscores the conserved role of these kinase pathways in mechanical regulation. It is important to note, that the study by Toyoda et al. (22) was conducted using AFM on adherent cells, whereas RT-FDC measures cells in suspension. In our case, we used the hemocyte-like *Drosophila* cell line *KC167*, which naturally grows in suspension and can thus be measured in its native mechanical state. Prior work has demonstrated that mechanical phenotypes can differ considerably between adherent and suspended cells, particularly in the context of cortical myosin II activity (62).

Notably, the *Drosophila* JAK homolog *hop* (*hopscotch*), identified in all three screens, had never been directly linked to cell mechanics. However, evidence from human studies suggests that the JAK signaling pathway may play a role in mechanical regulation (63, 64). The identification of the *JAK* homolog *hop* in all three screens provides additional support for this hypothesis and highlights the potential of genome-scale screening methods such as RT-FDC to uncover novel gene functions related to cell mechanics.

## CONCLUSION

We presented a genome-scale RNA interference (RNAi) knockdown screen investigating the roles of kinase and phosphatase genes in regulating single-cell mechanical phenotypes using Real-Time Fluorescence and Deformability Cytometry (RT-FDC). With 214 targeted genes, this study represents the largest deformability cytometry-based screen conducted to date. RT-FDC uniquely combines a high throughput of up to 100 cells/s with single-cell resolution, enabling comprehensive sampling of mechanical phenotype distributions within knockdown cell populations. Additionally, its capability to separately analyze interphase and mitotic cells allowed us to overcome the bias of population-wide analyses against softening and mitosis-specific knockdown effects, as well as avoiding apparent stiffening effects caused by underlying shifts in cell cycle distribution.

By identifying both known and novel mechanical regulators across diverse cellular processes, our results establish RT-FDC as a robust and versatile platform for large-scale mechanomic screening. Furthermore, the newly discovered mechanical regulators uncovered in our study, such as *PRL-1* and *hop*, provide valuable starting points for future investigations into genotype-phenotype relationships. Moving forward, mechanistic follow-up studies on both known and newly identified regulators will be essential for further unraveling how signaling pathways orchestrate structural determinants of cell mechanical phenotypes in both physiological and disease contexts. Among others, these might include gene overexpression, phenotype rescue or immunostaining, which could be facilitated by recent advances in protocols for staining cells in suspension (65, 66).

While our screen focused exclusively on cell stiffness quantified through a single Young’s modulus value, it is well-established that cells are viscoelastic, exhibiting both solid-like elastic and fluid-like viscous properties (67–69). Thus, representing cell mechanics by a single elastic modulus measured at sub-second time scales, as performed by RT-FDC, may overlook important viscous contributions and time-dependent deformation characteristics relevant to physiological conditions (see (15) for a detailed discussion). Recent advances in deformability cytometry address this limitation by leveraging the inherent modularity of the technique, allowing measurement modes to be adapted by simply exchanging the microfluidic chip design and adjusting the post-experimental data analysis pipeline. Examples include high-throughput viscoelastic phenotyping using hyperbolic microchannels (70) or the Microfluidic Microcirculation Mimetic (MMM), which analyzes cell traversal through microvascular-like constrictions (71). In addition, optical stretcher measurements may offer insights into cell stiffness on longer, second-scale timescales (11, 62), while atomic force microscopy (AFM) could facilitate direct comparisons with prior studies performed on adherent cells. The modularity of deformability cytometry facilitates straightforward adaptation of our screening approach to these advanced viscoelastic measurement techniques. We therefore anticipate future mechanomic screens incorporating viscoelastic phenotyping, enabling the identification of genetic regulators of cellular viscosity, and thus providing a more comprehensive understanding of cell mechanics beyond the current cytoskeleton-centered view.

## AUTHOR CONTRIBUTIONS

J.G. and B.B. designed the research and supervised the project. K.P., C.S, C.L., M.U. and M.K. performed the experiments. L.S. and K.P. analyzed the data. P.M. maintained analysis and data management software. L.S. and J.K. wrote the article.

## ACKNOWLEDGMENTS

This work is dedicated to Erich Sackmann, who has guided our understanding of mechanical cell properties and their structural determinants, especially the cytoskeleton. Our use of RT-FDC builds directly upon a legacy of innovation in biomechanical measurement techniques initiated by Erich Sackmann and continued by his students, including the PhD advisors of Jochen Guck and Jona Kayser. By building upon this foundation, we carry forward Erich Sackmann’s vision of gaining a comprehensive understanding of how cells actively regulate, maintain and adapt their mechanical state.

We want to thank Benedikt Hartmann, Felix Reichel, Ina Brauer, Nico Appold and Maximilian Eiche for helpful discussion regarding this study. Additional thanks to Nisarg Mehta and Raül González Garrido for their assistance during the recovery of archived measurement data, and to Eoghan O’Connell for help with data analysis. The authors acknowledge the financial support through the core funding of the Max Planck Society to J.G. and the Emmy Noether Programme of the German Research Foundation (455449456) to J.K.. B.B. received funding from Cancer Research UK: CRUK Programme Grant (C1529/A17343) and CRUK/EPSRC Multi-Disciplinary Project Award (C1529/A23335).

## DECLARATION OF GENERATIVE AI AND AI-ASSISTED TECHNOLOGIES IN THE WRITING PROCESS

During the preparation of this work the authors used ChatGPT-4o and o1 in order to improve readability and language. After using this tool, the authors reviewed and edited the content as needed and take full responsibility for the content of the publication.

## DECLARATION OF INTERESTS

Jochen Guck, Martin Kräter and Paul Müller are cofounders and shareholders of the company Rivercyte GmbH, which commercializes deformability cytometry devices, consumables, and software for clinical applications.

## SUPPLEMENTAL MATERIAL

**Table S1** List of all targeted genes, their dsRNA strands used for RNAi knockdown and their sample IDs used during measurement and analysis of the screen.

**Table S2** List of all mechanical hits identified by the cell cycle-resolved analysis. For each gene, its Young’s modulus change, significance and error margins for each analysis (cell cycle-resolved analysis, full population analysis and full population analysis after mitotic downsampling) are recorded. In addition, mitotic fraction change upon knockdown is recorded for each gene.

**Table S3** List of all mechanical hits identified exclusively by the full population analysis. For each gene, its full population Young’s modulus change, significance and error margins before and after mitotic downsampling are recorded. In addition, mitotic fraction change upon knockdown is recorded for each gene.

## Notes

https://owncloud.gwdg.de/index.php/s/cwsnBzHnIdqtRFS

### Competing Interest Statement

Jochen Guck, Martin Kr\"ater and Paul M\"uller are cofounders and shareholders of the company Rivercyte GmbH, which commercializes deformability cytometry devices, consumables, and software for clinical applications.

### Summary of Updates

Results section reorganized to present the cell cycle-resolved analysis as the central analytical approach from the beginning to avoid apparent self-contradictions; analysis and discussion of full-population-only hits moved to Supplemental Materials; Figure 1 and 2 combined; author list updated; declaration of interests updated.

